# Microstimulation of sensory cortex engages natural sensory representations

**DOI:** 10.1101/2022.11.09.515803

**Authors:** Ravi Pancholi, Andrew Sun-Yan, Simon Peron

**Author notes:** Lead Contact: Simon Peron.

## Abstract

Cortical activity patterns occupy a small subset of possible network states. If this is due to intrinsic network properties, microstimulation of cortex should evoke activity patterns resembling those observed during natural sensory input. Here, we use optical microstimulation of virally transfected layer 2/3 pyramidal neurons in mouse primary vibrissal somatosensory cortex to compare artificially evoked activity with natural activity evoked by whisker touch and movement (‘whisking’). We find that photostimulation engages touch but not whisking responsive neurons more than expected by chance. Neurons that respond to photostimulation and touch or to touch alone exhibit higher spontaneous pairwise correlations than purely photoresponsive neurons. Exposure to several days of simultaneous touch and optogenetic stimulation raises both overlap and spontaneous activity correlations among touch and photoresponsive neurons. We thus find that cortical microstimulation engages existing cortical representations and that repeated co-presentation of natural and artificial stimulation enhances this effect.

## INTRODUCTION

Cortical activity occupies a small subspace of possible activity patterns (Avitan and Stringer, 2022; Chung and Abbott, 2021; Gallego et al., 2017; Jazayeri and Afraz, 2017; Okun et al., 2015; Tsodyks et al., 1999). For instance, spontaneous activity often resembles activity evoked by natural stimuli, an observation attributed to intrinsic constraints imposed by cortical connectivity and variable physiology among individual neurons (Kenet et al., 2003; Luczak et al., 2009). Intrinsic constraints on evoked activity patterns may explain why microstimulation of sensory cortex can evoke naturalistic sensory hallucinations (Penfield and Boldery, 1937) while motor cortical stimulation can drive complex, naturalistic movements (Graziano et al., 2002; Penfield and Boldery, 1937).

Cortical networks are organized in a highly specific manner. Reciprocal connectivity among small groups of pyramidal neurons far exceeds what is expected by chance (Perin et al., 2011; Song et al., 2005). Synaptic connectivity is also heterogeneous, with a few very strong synapses among a much larger pool of weak synapses (Lefort et al., 2009). In layer (L) 2/3 of mouse primary visual cortex, neurons with similar tuning are far more likely to be interconnected (Cossell et al., 2015; Ko et al., 2011; Lee et al., 2016; Wertz et al., 2015), as are cortical neurons that receive common input from thalamus (Morgenstern et al., 2016) or other cortical layers (Yoshimura et al., 2005). Taken together, these observations suggest that cortex consists of small, highly interconnected subnetworks in a sea of weaker connectivity. Such structured connectivity should greatly constrain the response to random cortical stimulation, favoring pattern completion (Douglas and Martin, 2007) that engages these highly interconnected subnetworks.

Cortical microstimulation has been used extensively in the study of perception, with both somatosensory (Romo et al., 1998) and visual cortical (Afraz et al., 2006; Salzman et al., 1990) stimulation biasing perception. However, due to the difficulties of recording during microstimulation, it remains unclear whether microstimulation engages activity patterns similar to those evoked by natural stimuli, thereby accounting for perceptual biasing. Cellular resolution two-photon optogenetic experiments, which explicitly stimulate a handful of opsin-expressing neurons with known tuning while recording calcium activity, reveal that perception can be biased specifically toward the tuning of the stimulated neurons and that effective pattern completion in the form of observable network activation predicts perceptual effect (Carrillo-Reid et al., 2019; Marshel et al., 2019). Because optical microstimulation is compatible with concurrent recording, it opens the door to addressing the question of whether intrinsic network features sufficiently constrain cortical activity to ensure that evoked activity will resemble natural cortical responses even when neurons with specific receptive fields are not explicitly targeted.

Here, we combine optical microstimulation with volumetric two-photon microscopy (Peron et al., 2015a; Peron et al., 2015b) of L2/3 of primary vibrissal somatosensory cortex (vS1) to ask whether directly driven cortical activity engages the same neural populations as naturalistic stimuli. First, we assess the responsiveness of vS1 neurons to vibrissal touch and movement (‘whisking’). Next, we assess the response of the same neurons to direct optogenetic stimulation. Because touch but not whisking neurons are recurrently coupled (Peron et al., 2020), only touch networks should engage in pattern completion characteristic of recurrent networks (Douglas and Martin, 2007). Thus, we expect that optogenetic activation of vS1 should engage touch, but not whisking networks, a prediction borne out by our results.

We next sought to ask whether this effect could be enhanced by repeated co-stimulation of the natural and photoresponsive populations. Hebbian plasticity should drive elevated connectivity among co-stimulated neurons (Clopath et al., 2010; Litwin-Kumar and Doiron, 2014), especially given the spike timing dependent plasticity rule governing L2/3 pyramidal neurons in vS1 (Banerjee et al., 2014). We therefore expose mice to an induction paradigm in which optical microstimulation of vS1 was paired with concurrent presentation of a vibrissal touch stimulus. We find that this enhances the overlap among the photoresponsive and touch responsive populations. Our work suggests that intrinsic features of cortical circuitry bias activity toward response patterns observed during naturalistic stimulation, and that this biasing can be enhanced through repeated co-presentation of natural and artificial stimuli.

## RESULTS

### Recording photostimulus-evoked and vibrissal sensory activity in opsin-transfected sensory cortex

To assess the relationship between artificially evoked neural activity and natural sensory responses, we measured cortical activity as mice were presented with direct cortical stimulation or natural vibrissal stimuli. Transgenic mice expressing GCaMP6s in cortical excitatory neurons (Slc17a7-Cre X Ai162) (Daigle et al., 2018) were virally transfected with the soma-restricted opsin ChRmine (AAV-8-CaMKIIa-ChRmine-mScarlet-Kv2.1-WPRE) (Marshel *et al*., 2019) in layer (L) 2/3 of primary vibrissal somatosensory cortex (vS1) and implanted with a cranial window (**Figure 1A**). Following recovery, mice were trimmed to a single row of whiskers (C row) whose barrel columns were localized using widefield two-photon calcium imaging (Methods; **Figure S1**). Screening following the onset of viral expression confirmed that opsin was restricted to vS1 and overlapped with the C row barrel columns. Animals were trimmed to the two whiskers whose barrel columns best overlapped the opsin-expressing area. A miniature light-emitting diode (LED) was affixed to the cranial window adjacent to the site of opsin expression for optogenetic stimulation.

**Figure 1.**
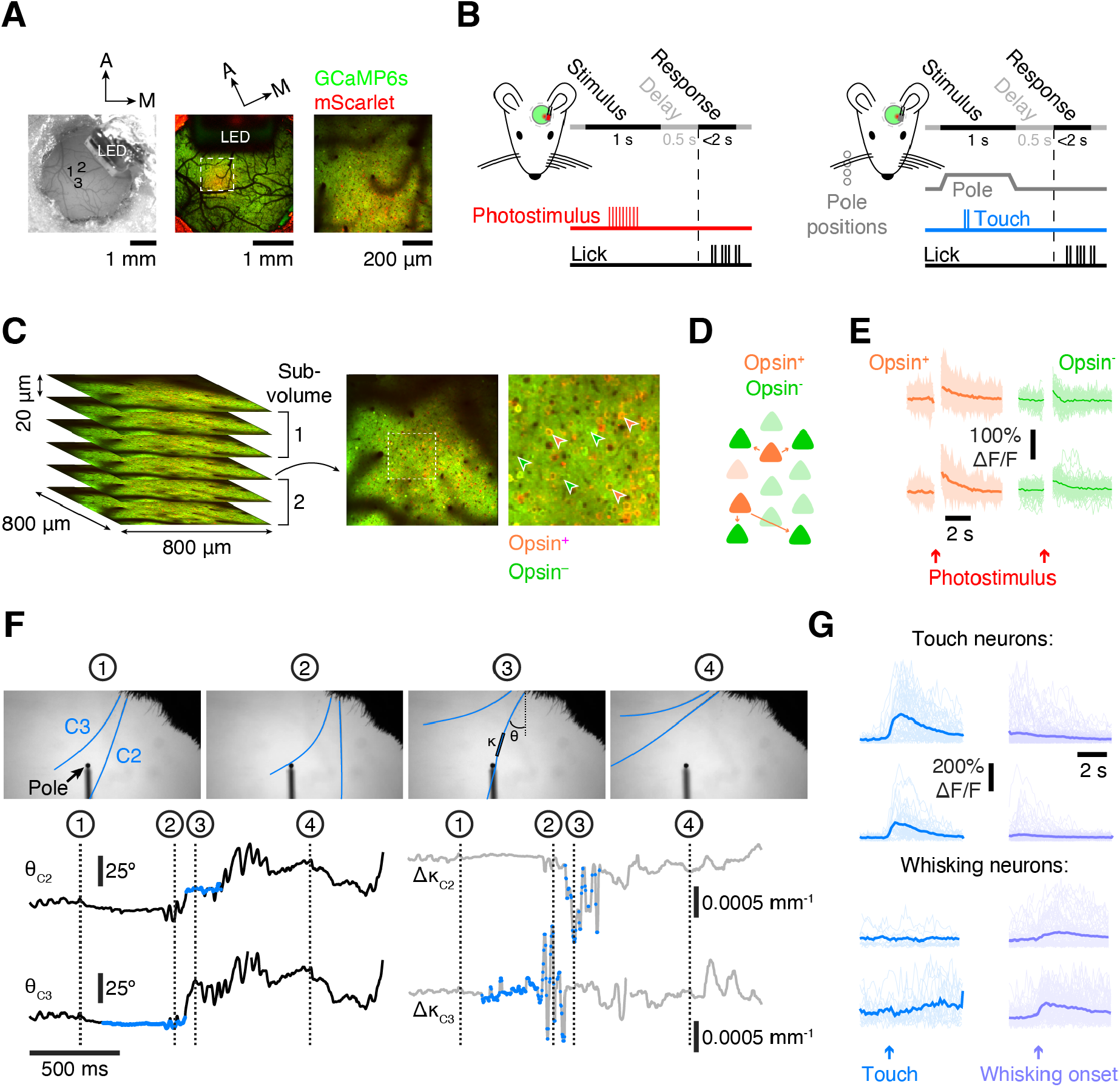
Neurons in L2/3 of vS1 show photostimulus, touch, and whisking evoked activity. (A) Opsin expression in barrel cortex. Left, widefield view of cranial window showing viral injection sites (numbers) and LED; center, widefield two-photon image of cranial window after onset of opsin expression (green, GCaMP6s fluorescence; red, mScarlet fluorescence); right, higher magnification two-photon image. (B) Left, trial structure during a rewarded photostimulation trial. Right, trial structure during a rewarded vibrissal touch trial. (C) Volumetric two-photon imaging. Subvolumes consist of 3 planes imaged simultaneously at 7 Hz. Left, six planes (800-by-800 μm, 20 μm inter-plane distance, 2 subvolumes); right, example plane with example opsin-expressing (orange) and opsin non-expressing (green) neurons. (D) Schematic showing the expected local propagation of activity from opsin-expressing (orange) to opsin non-expressing (green) neurons. (E) Photostimulation-evoked ΔF/F traces for 4 neurons. Light, individual trials; dark, mean. Orange, opsin-expressing; green, opsin non-expressing. A PMT shutter prevented acquisition during photostimulation (Methods). (F) Whisker videography in an example trial. Top, example frames; bottom, whisker angle (θ, black), and change in whisker curvature (Δκ, grey, see Methods), with touch periods indicated with colored circles. (G) ΔF/F traces for 4 neurons aligned to whisker touch (left column) and to whisk onset (right column). Grey, individual trials; black, mean. Top two neurons, touch neurons; bottom two neurons, whisking neurons.

To assess the response to direct cortical activation, we stimulated the opsin expressing neurons with the LED (9 pulses, 20 Hz, 5 ms; **Figure 1B**). In some mice, photostimulation was followed by water reward on a subset of trials; in others, no reward accompanied photostimulation (Methods; **Table S1**). The touch response was assessed in a separate session by presenting mice with a pole that appeared in one of several positions. To encourage active whisking, were given a water reward after pole withdrawal on a subset of trials (Methods). In both touch and photostimulation sessions, trials began with a 1s stimulus epoch. On water rewarded trials, this was followed by a short delay period (500 ms), after which an auditory cue alerted mice that they could lick for a reward during the response epoch (<2 s). In both cases, an interstimulus interval of 10-15 s was employed.

We recorded neural activity during stimulation using volumetric two-photon calcium imaging. On each trial, we simultaneously tracked neurons distributed among 3 imaging planes (800-by-800 μm, 20 μm spacing, one ‘subvolume’) at 7 Hz (**Figure 1C**). We alternated between two subvolumes throughout each session, switching subvolumes every ∼100-150 trials. Imaged neurons were separated into opsin-expressing and opsin non-expressing based on the presence of the opsin-conjugated fluorophore, mScarlet (Methods). We found photostimulus-evoked responses among neurons in both populations, with opsin non-expressing neurons presumably excited by opsin-expressing neurons in a feedforward manner (**Figure 1D, E**).

To precisely determine times of touch and measure whisker kinematics, we recorded whisker movement using high-speed whisker videography (**Figure 1F**). Whisker position was defined as the azimuthal angle (θ) of the whisker with the face. Tactile input was measured by the touch-induced change in whisker curvature (Δκ), which is proportional to the force acting on the whisker follicle and correlated with the activity of contact-responsive neurons in vS1 (Severson et al., 2017). We found neurons responsive to both whisker movement (‘whisking neurons’) and object contact (‘touch neurons’) within the recorded population (**Figure 1G**). Thus, in a volume of vS1 transfected with an opsin, both photostimulation and natural touch evoke robust neural responses.

### Photostimulation and touch responsive populations overlap more than expected by chance

Does optical microstimulation of sensory cortex disproportionately engage natural sensory representations? To assess this, we compared photostimulation and touch responsiveness across neurons in vS1, comparing touch-evoked activity with photostimulation-evoked activity from two separate imaging sessions (Methods). Touch and photostimulation each engaged a substantial population of neurons (**Figure 2A**), with a subset of neurons responding to both stimuli (**Figure 2B**). Neurons that responded to both stimuli tended to respond less strongly than the neurons most responsive to a given stimulus (**Figure 2C**).

**Figure 2.**
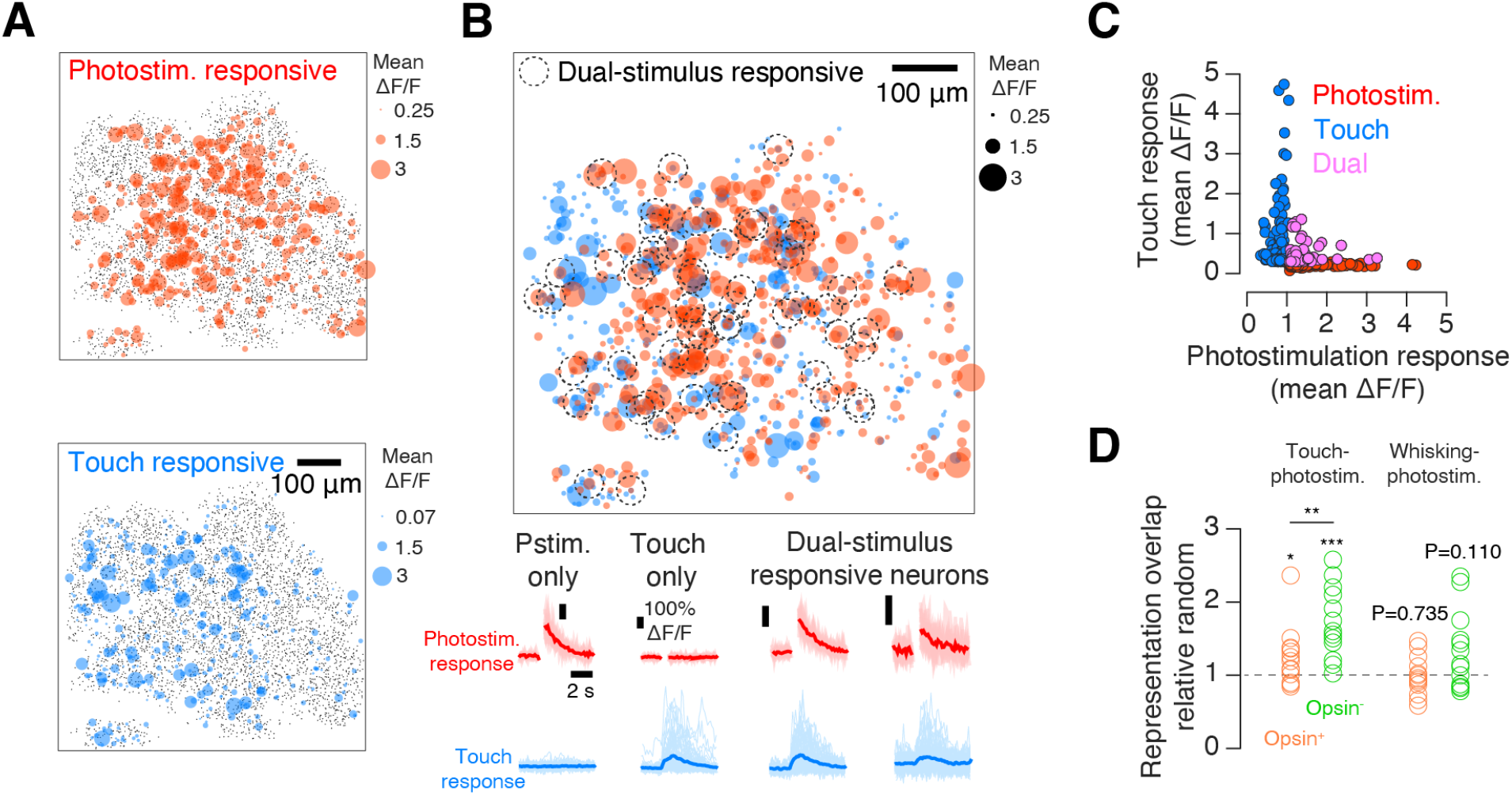
Overlap between touch responsive and photoresponsive populations. (A) Photoresponsive (top, red) and touch responsive (bottom, blue) populations in an example mouse. Neurons are collapsed across six planes spaced 20 μm apart. Colored circles show the mean touch (blue) or photostimulation (red) evoked ΔF/F across stimulus presentations. Black dots indicate neurons that did not belong to the top 10% of responders (Methods). (B) Top, overlay of the maps in (A), with non-responsive neurons removed. Neurons belonging to both representations are marked with a black dotted circle. Bottom, stimulus-aligned photostimulation (red, top) and touch (blue, bottom) ΔF/F responses of four example neurons. Dark color, mean response; light color, individual responses. (C) Relationship between mean ΔF/F touch and photostimulation responses for all responsive neurons in an example mouse. Blue, neurons that belong only to the top 10% of touch responders; red, neurons that belong only to the top 10% of photostimulation responders; magenta, neurons that belong to both groups (‘dual’). (D) Overlap between vibrissal and photostimulation responsive populations relative to chance for opsin-expressing (orange) and opsin non-expressing (green) neurons across mice (N=13). P-values are given for the Wilcoxon signed rank test assessing whether the median is different than 1 (chance); P-values between opsin-expressing and opsin non-expressing populations are for the paired Wilcoxon signed rank test. Left, overlap between touch responsive and photoresponsive population; right, overlap between whisking responsive and photoresponsive population.

To quantify the similarity between the two network responses, we asked whether the number of neurons responding to both stimuli was greater than, equal to, or less than the number expected by chance. Specifically, we computed the number of neurons that would be expected to belong to both representations assuming random draws with replacement from a given source population (Methods). Dividing the actual number of dual-representation neurons by this number, we obtained a randomness-normalized overlap measure, with values equal to one implying no deviation from randomness, values less than one implying lower-than-expected overlap, and values greater than one implying greater-than-expected overlap. For touch and photostimulus responsive neurons, we measured a normalized overlap of 1.25 ± 0.40 (mean ± S.D., N=13 mice) among opsin expressing neurons (**Figure 2D**), a value significantly greater than one (P=0.027, Wilcoxon signed rank test assessing whether the ratio was equal to 1). Thus, opsin expressing neurons that likely respond directly to optical microstimulation show an elevated likelihood of being touch responsive as well.

Because we directly stimulate only opsin-expressing neurons, we reasoned that overlap should be greater in photoresponsive opsin non-expressing neurons, as these neurons are activated indirectly due to propagation of activity through the network and should therefore be more susceptible to intrinsic constraints on response. Consistent with this, the normalized touch-photostimulation overlap was 1.73 ± 0.48 among opsin non-expressing neurons, which was significantly greater than one (P<0.001) and exceeded the overlap observed in opsin-expressing neurons (P=0.005, Wilcoxon signed rank test comparing opsin-expressing overlap to opsin non-expressing overlap, paired by animal).

To assess the degree to which the photostimulation response among opsin-expressing neurons was under experimental control, we examined how well opsin levels could predict the photostimulation response. We found that the level of opsin expression, quantified using ‘redness’ (**Figure S2A**; Methods), a measure of the amount of mScarlet (Bindels et al., 2017) co-expressed with the opsin ChRmine, was correlated to response amplitude among opsin-expressing neurons (**Figure S2B**). Across animals, this correlation was significantly greater than zero (**Figure S2C**, 0.15 ± 0.08 ; P<0.001, Wilcoxon signed rank test assessing if the cross-animal median differs from zero). Thus, opsin expression levels explain some, but not all, of the evoked neural response.

Given that touch but not whisking neurons are recurrently coupled (Peron *et al*., 2020), if overlap is a consequence of pattern completion due to recurrence, whisking neurons should not overlap with photoresponsive neurons more than expected by chance. For opsin-expressing neurons, we measured a whisking-photoresponsive overlap ratio of 0.99 ± 0.26, a value not significantly different from 1 (P=0.735). Whisking-photoresponsive overlap for opsin non-expressing neurons was also not significantly different from 1 (1.32 ± 0.53; P=0.110). Thus, whereas the touch responsive population overlapped more than expected by chance with the photoresponsive population, the whisking population did not.

We thus find that dual-representation touch-photoresponsive neurons are more common than expected by chance, whereas dual-representation whisking-photoresponsive neuron are not.

### Spontaneous activity correlations are higher among dual-representation neurons

Touch and photoresponsive populations may overlap disproportionately because photostimulation of dual-representation neurons engages recurrence-based pattern completion among touch neurons. In this situation, correlations among neurons with elevated coupling should be higher (Cossell *et al*., 2015; Ko *et al*., 2011). We therefore examined pairwise correlations within photoresponsive, touch responsive and dual-stimulus responsive populations. Because correlations during photostimulation would be at least partly driven by direct optogenetic activation, we restricted our analysis to ‘spontaneous’ activity during the interstimulus epoch on touch-only imaging sessions (Methods; **Figure 3A**).

**Figure 3.**
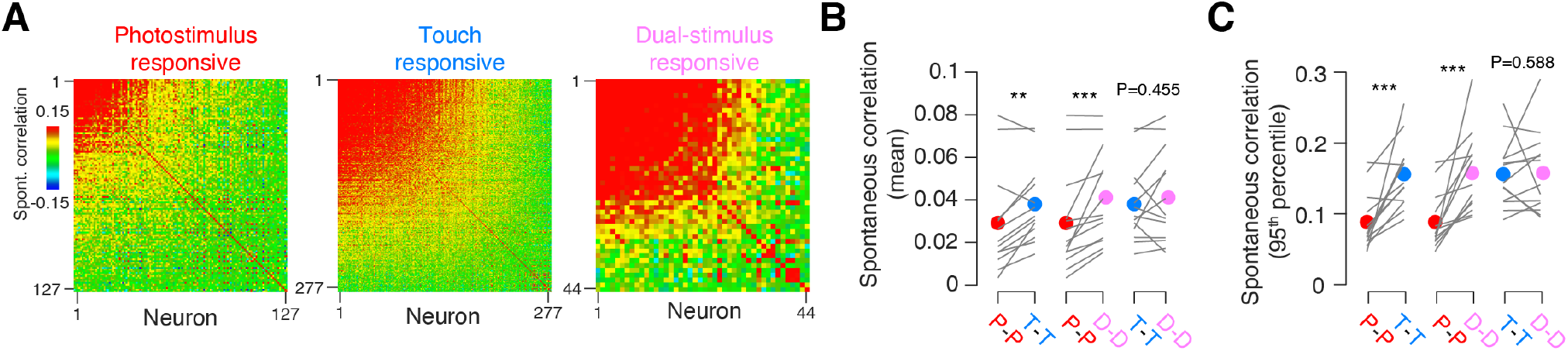
Spontaneous activity correlations within different populations. (A) Pairwise correlations during the inter-stimulus (‘spontaneous’) period in an example mouse. Left, photostimulus responsive neurons. Middle, touch responsive neurons. Right, dual-stimulus responsive neurons. Neurons are sorted by their mean correlation to the other neurons within the population. (B) Mean spontaneous pairwise correlations across mice (N=13) for photoresponsive (red), touch responsive (blue), and dual-stimulus responsive (magenta) neurons. P-value provided Wilcoxon signed rank test comparing two populations, paired by animal. ***, P < 0.001; **, P < 0.01 ; *, P < 0.05. (C) As in (B), but for 95^th^ percentile of spontaneous pairwise correlations.

Spontaneous pairwise correlations among neurons that were only photoresponsive were lower than among neurons responsive only to touch (photoresponsive: 0.03 ± 0.02, touch: ± 0.02, mean ± S.D., N=13 mice; P=0.008, Wilcoxon signed rank test comparing within-photoresponsive and within-touch responsive pairwise correlations; **Figure 3B**). However, the dual-stimulus responsive population showed within-population correlations comparable to the touch population (0.04 ± 0.02; P=0.455), and greater correlations than the photoresponsive population (P=0.001). This effect was especially pronounced among the most correlated cells: the cross-animal mean of 95^th^ percentiles of pairwise correlations among photoresponsive neurons was 0.09 ± 0.04 (**Figure 3C**), substantially less than correlations among touch responsive neurons (0.15 ± 0.05; P=0.001). Dual-representation neurons, however, showed pairwise correlations indistinguishable from touch neurons (0.16 ± 0.06; P=0.588), and substantially greater than photoresponsive neurons (P<0.001). Thus, neurons responsive to both touch and photostimulation show spontaneous activity correlations comparable to those in touch neurons, whereas neurons responsive only to photostimulation show substantially lower spontaneous pairwise correlations.

### Combined touch-photostimulation stimulation drives increased population overlap

Can overlap be increased? The plasticity rules governing vS1 L2/3 (Banerjee *et al*., 2014) suggest that two populations should experience enhanced connectivity due to Hebbian plasticity if neurons are co-activated in a short enough time window. We therefore presented mice with a proximal pole and, on the same trials, photostimulated opsin-expressing neurons with our 9-pulse stimulus (Methods; **Figure 4A**). Mice were stimulated in this manner over 10 days. A subset of trials featured photostimulation with no pole, allowing us to evaluate the photostimulus-only response, which we did using imaging on the first and final days of this induction protocol. A separate touch-only session (Methods) preceded the first induction session and another followed the last induction session; these sessions were used to assess touch responsiveness. Vibrissal kinematics did not exhibit systematic differences across pre- and post-induction touch sessions (**Figure S3**).

**Figure 4.**
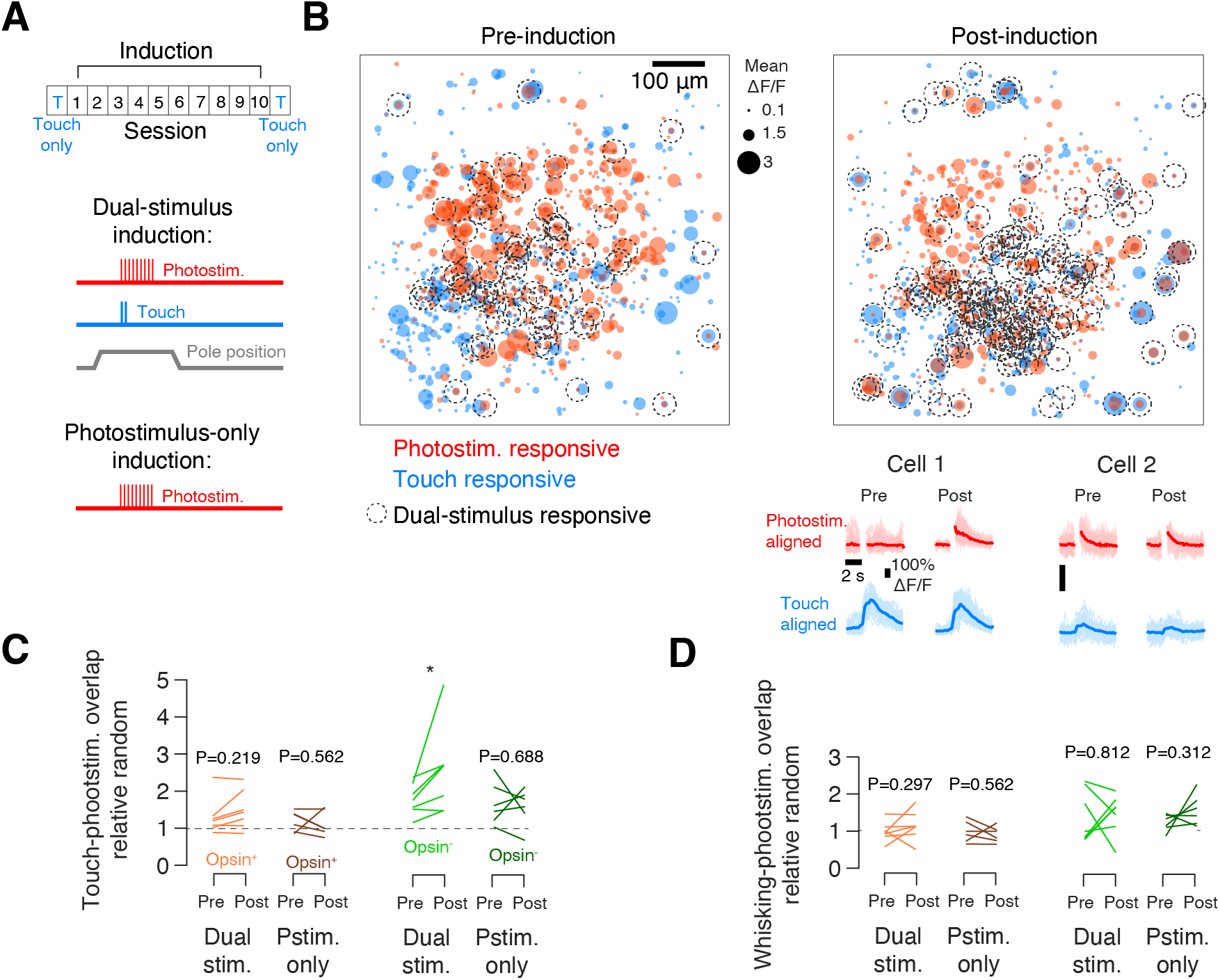
Change in overlap between touch and photoresponsive populations following induction. (A) Timeline of the induction experiment. Top, breakdown of sessions in induction experiment. Ten session induction block is preceded and followed by touch-only sessions (Methods). Middle, dual-stimulus induction features photostimulation (9 pulses; Methods) during a period of pole accessibility. Bottom, photostimulation-only induction does not include pole presentation. (B) Maps showing the photoresponsive (red) and touch responsive (blue) populations in an example mouse before (left) and after (right) dual-stimulus induction. Colored circles show the mean touch (blue) or photostimulus (red) evoked ΔF/F across stimulus presentations. Neurons belonging to both representations are circled with a dotted line. Bottom, response of two example neurons prior to (left) and after (right) induction. (C) Touch-photostimulation overlap among opsin-expressing (orange) and opsin non-expressing (green) populations before and after induction. Mice were exposed to dual-stimulus induction (lighter color, N=7 mice) or photostimulation-only induction (darker color, N=6 mice). P-values provided for signed rank test comparing response before and after induction. ***, P < 0.001; **, P < 0.01 ; *, P < 0.05. (D) As in (C), but for whisking-photoresponsive overlap.

Induction increased overlap between the two populations (**Figure 4B**). Among opsin non-expressing neurons, overlap increased from 1.78 ± 0.42 to 2.54 ± 1.17 following dual-stimulus induction (**Figure 4C**; P=0.031, Wilcoxon signed rank test comparing pre- and post-induction overlap paired by animal; N=7 mice). This increase was not observed in the opsin-expressing population, in which the overlap between touch and photoresponsive populations was 1.31 ± 0.49 pre-induction and 1.49 ± 0.52 post-induction (P=0.219). Thus, repeated co-presentation of touch and photostimulation resulted in increased representational overlap among opsin non-expressing neurons.

In a second group of mice (N=6), induction consisted of 10 days of repeated optical microstimulation with no touch or reward (**Figure 4A**). In these animals, overlap did not increase in either opsin expressing or opsin non-expressing populations (opsin expressing overlap: 1.19 ± 0.28 to 1.12 ± 0.34, P=0.562, N=6 mice; opsin non-expressing: 1.67 ± 0.58 to 1.57 ± 0.51, P=0.688). Moreover, overlap among whisking and photoresponsive populations did not increase in either population following both dual-stimulus induction and photostimulation-only induction (**Figure 4D**). Thus, dual-stimulus induction increases overlap between touch and photoresponsive populations in opsin non-expressing neurons, whereas photostimulation-only induction does not.

We next examined the properties of post-induction dual-responsive neurons. In contrast to pre-induction, post-induction dual-stimulus responsive neurons had high responses to both photostimulation and touch (**Figure 5A**). Did overlap increase due to more touch cells becoming photoresponsive, more photoresponsive cells becoming touch responsive, or both? Among mice exposed to dual-stimulus induction, the fraction of opsin-expressing photoresponsive neurons that also responded to touch did not change (from 0.13 ± 0.05 to 0.15 ± 0.07; P=0.375, Wilcoxon signed rank test comparing pre- and post-induction fraction; **Figure 5B**). Similarly, no change in touch response fraction among photoresponsive neurons was seen in mice exposed to photostimulation-only induction (from 0.11 ± 0.04 to 0.10 ± 0.05, P=0.562). Among opsin non-expressing neurons in dual-stimulus induction mice, however, the touch responsive fraction increased from 0.18 ± 0.05 to 0.26 ± 0.11 (P=0.016); photostimulation-only induction did not drive any change among these neurons (from 0.17 ± 0.06 to 0.16 ± 0.05, P=0.688). The fraction of touch neurons that were also photoresponsive also only changed among opsin non-expressing neurons in dual-stimulus induction mice (from 0.10 ± 0.05 to 0.15 ± 0.05; P=0.016; **Figure 5C**). No change in the fraction of touch responsive neurons with a photostimulus response took place among opsin-expressing neurons in dual-stimulus induction mice (from 0.31 ± 0.11 to 0.31 ± 0.14, P=0.938) or photostimulation-only induction mice (from 0.36 ± 0.13 to 0.35 ± 0.21, P=0.438), or among opsin non-expressing neurons in photostimulation-only induction mice (from 0.07 ± 0.03 to 0.07 ± 0.02, P=1). Thus, dual-stimulus induction increases both the fraction of touch cells that are photoresponsive and photoresponsive cells that are touch responsive, but only among opsin non-expressing neurons.

**Figure 5.**
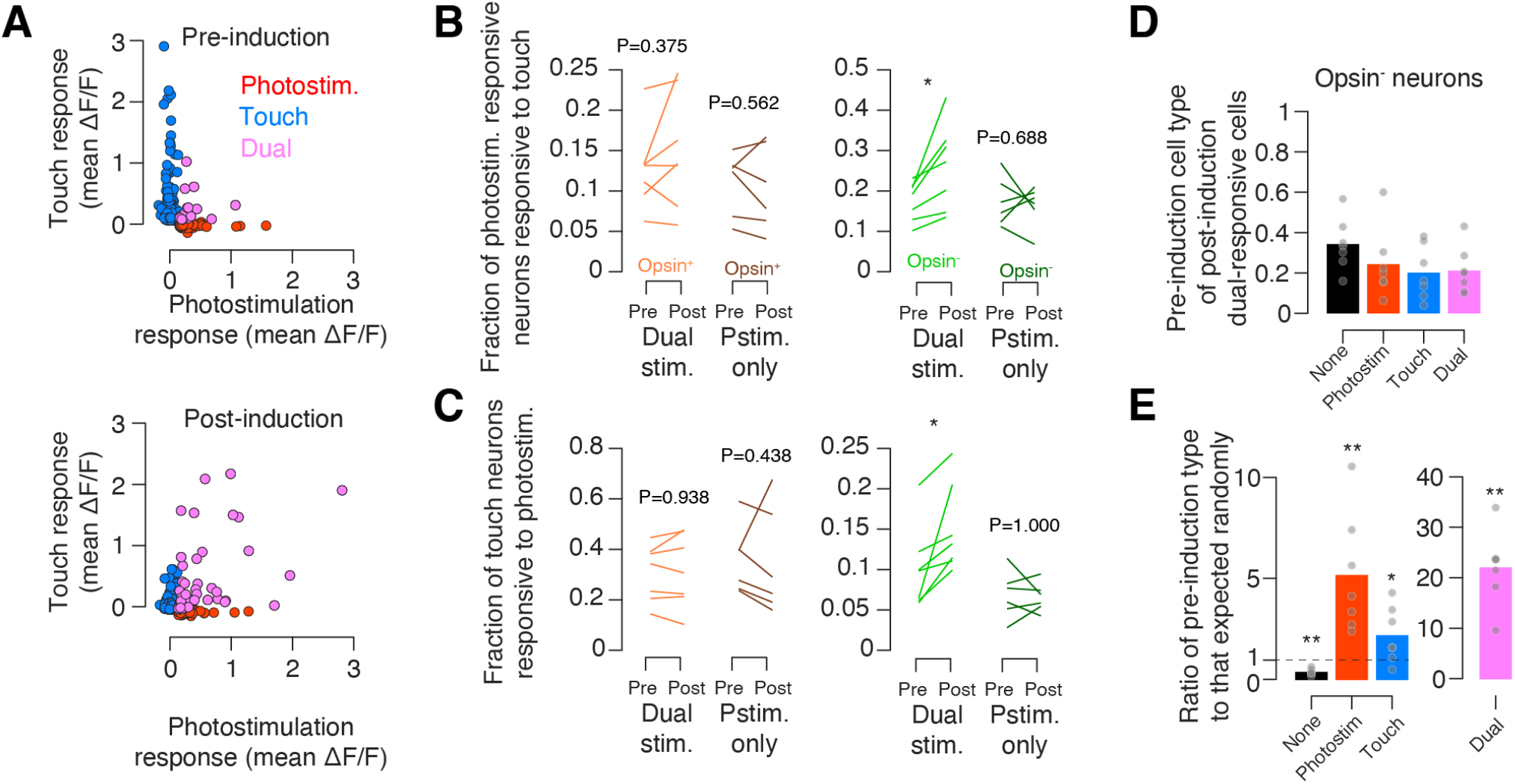
Basis of increased touch-photostimulation population overlap following induction. (A) Change in relationship between mean ΔF/F response to touch and photostimulation for all responsive neurons in an example mouse. Blue, neurons that belong only to the top 10% of touch responders; red, neurons that belong only to the top 10% of photostimulation responders; magenta, neurons that belong to both groups (‘dual’). Top, relationship prior to induction. Bottom, relationship following induction. (B) Fraction of photoresponsive neurons that also respond to touch before and after induction. Orange, opsin-expressing neurons; green, opsin non-expressing. Mice were exposed to dual-stimulus induction (lighter color, N=7 mice) or photostimulation-only induction (darker color, N=6 mice). P-values provided for signed rank test comparing fraction before and after induction. ***, P < 0.001; **, P < 0.01 ; *, P < 0.05. (C) As in (B), but showing the fraction of touch responsive neurons that also respond to photostimulation. (D) Fraction of neurons that were responsive to both photostimulation and touch following dual-stimulus induction having a given cell type prior to induction. Dots, individual animals. Bars, mean (N=7 mice). (E) As in (D), but normalized to the size of each population on the post-induction day (i.e., what would be expected if neurons changed type randomly). P-values for signed rank test comparing the observed values to a distribution with median 0. ***, P < 0.001; **, P < 0.01; *, P < 0.05.

We next asked which populations contributed most to the post-induction dual-representation population. We restricted our analysis to opsin negative neurons in dual-stimulus induction mice, as only these neurons exhibited a change in overlap following induction. Of these neurons, 0.21 ± 0.12 (fraction; N=7 mice) were dual-stimulus responsive prior to induction, 0.24 ± 0.17 were photoresponsive only, 0.20 ± 0.13 were touch responsive only, and 0.21 ± 0.12 did not respond to either stimulus on the first day (**Figure 5D**). Thus, post-induction dual-stimulus responsive neurons were roughly equally drawn from all four possible sources. When normalized to the size of the pre-induction populations, however, previously responsive neurons were far more likely than chance to become dual-stimulus responsive: touch neurons were 2.22 ± 1.37 times more likely than chance to become dual-stimulus responsive (P=0.047, Wilcoxon signed rank test determining if median ratio was equal to 1; **Figure 5E**), photoresponsive neurons were 5.20 ± 2.95 times more likely than chance to become dual-stimulus responsive (P=0.016), and dual-stimulus responsive neurons were 22.15 ± 7.29 times more likely than chance to remain dual-stimulus responsive (P=0.016). Moreover, non-responsive neurons were significantly less likely than chance to become dual-stimulus responsive (0.40 ± 0.15; P=0.016). Thus, neurons that were initially responsive, especially dual-stimulus responsive neurons, were the most likely to become dual-stimulus responsive following induction.

### Combined touch-photostimulation training sparsens stimulus response

Did overlap and pairwise correlations between touch and photoresponsive populations rise because new neurons emerged that were responsive to both stimuli, or because induction reduced responsiveness among neurons that did not respond to both stimuli? To assess this, we examined how the aggregate responsiveness changed to both photostimulation and touch following dual-stimulus induction.

Following induction, the responsiveness and membership of both populations changed (**Figure 6A**). The mean ΔF/F response to photostimulation among photoresponsive opsin-expressing neurons (defined as the top 10% of responders; Methods), declined (**Figure 6B**) from 0.65 ± 0.19 to 0.39 ± 0.10 (mean ± S.D.; N=7 mice; P=0.004, Wilcoxon signed rank test comparing pre- and post-induction response, paired by animal). Opsin-expressing neurons in mice exposed to photostimulation alone did not show a change in their photostimulus response (0.75 ± 0.32 to 0.59 ± 0.25, N=6 mice; P=0.394), nor did opsin non-expressing neurons in dual-stimulus induction animals (0.59 ± 0.25 to 0.43 ± 0.11; P=0.259) or photostimulation-only induction animals (0.64 ± 0.20 to 0.51 ± 0.14; P=0.240). The mean touch response among touch responsive opsin-expressing neurons also declined (**Figure 6C**), from a mean ΔF/F of 0.19 ± 0.08 to 0.12 ± 0.03 (P=0.017, Wilcoxon signed rank test comparing pre- and post-induction response, paired by animal). Again, opsin-expressing neurons in mice exposed to photostimulation-only induction did not show a significant change in touch response (0.14 ± 0.05 to 0.13 ± 0.02; P=0.485), nor did opsin non-expressing touch neurons in dual-stimulus induction animals (0.20 ± 0.07 to 0.17 ± 0.04; P=0.456) or photostimulation-only animals (0.18 ± 0.06 to 0.17 ± 0.02; P=0.310). Thus, both touch and photostimulus responses declined among opsin-expressing neurons in dual-stimulus induction animals.

**Figure 6.**
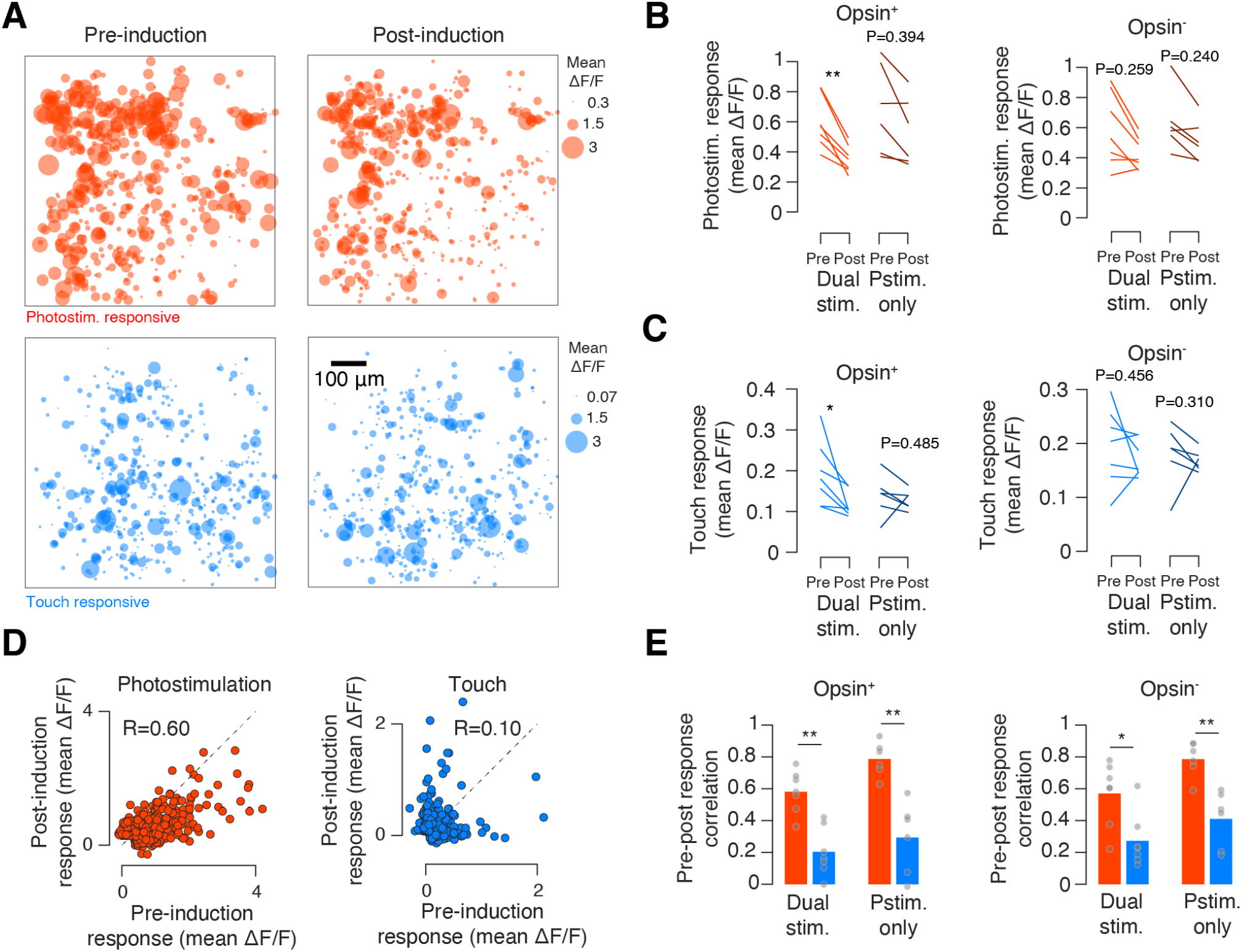
Change in touch and photoresponsive populations following induction. (A) Photostimulation (top, red) and touch-responsive (bottom, blue) populations in an example mouse before (left) and after (right) dual-stimulus induction. Neurons are collapsed across six planes spaced 20 μm apart. Colored balls show the mean touch (blue) or photostimulation (red) evoked ΔF/F across stimulus presentations. (B) Change in mean stimulus-evoked ΔF/F among the top 10% of most photoresponsive neurons following induction. Left, opsin-expressing; right, opsin non-expressing. In each case, we examined mice trained on dual-stimulus induction (N=7; lighter color), and photostimulation-only induction (N=6; darker color). P-values provided for signed rank test comparing response before and after induction. ***, P < 0.001; **, P < 0.01 ; *, P < 0.05. (C) As in (B), but for touch responsiveness. (D) Stimulus response (mean evoked ΔF/F) before and after dual-stimulus induction. Left (red), photostimulation response among all responsive neurons before and after induction in an example mouse. Right (blue), touch response before and after induction. (E) Correlation of pre- and post-induction mean evoked ΔF/F for photostimulation responsive (red) and touch responsive (blue) populations. Left, opsin-expressing; right, opsin non-expressing neurons. ***, P < 0.001; **, P < 0.01 ; *, P < 0.05; P-value for Wilcoxon signed rank test comparing touch and photostimulus responsive populations.

Responsiveness for individual members of a sensory cortical representation tends to change over time in the face of constant input, a process known as ‘representational drift’ (Marks and Goard, 2021; Schoonover et al., 2021). To compare drift across populations, we computed the mean stimulus-evoked ΔF/F for each neuron pre- and post-induction for a given stimulus. We then measured the Pearson correlation between these two vectors. The photoresponsive population was consistently more stable than the touch responsive population (**Figure 6D**). Among animals exposed to dual-stimulus induction (N=7 mice), the mean correlation between pre- and post-induction photostimulation responses among photoresponsive neurons (**Figure 6E**; R_photostim_=0.58 ± 0.13) was higher than the correlation of touch responses among touch neurons (R_touch_=0.20 ± 0.15; P=0.002, Wilcoxon signed rank test comparing touch and photoresponsive correlations, paired by animal). Photoresponsive populations were more stable than touch populations in opsin-expressing neurons of mice exposed to photostimulus-only induction (R_photostim_=0.79 ± 0.11, R_touch_=0.29 ± 0.22 P=0.002), and opsin non-expressing neurons of mice exposed to both dual-stimulus induction (R_photostim_=0.57 ± 0.20, R_touch_=0.27 ± 0.17; P=0.026) and photostimulus-only induction (R_photostim_=0.79 ± 0.11, R_touch_=0.41 ± 0.18; P=0.002). Thus, in all cases, photoresponsive populations were more stable than touch responsive populations.

Were the observed declines in responsiveness specific to the touch and photostimulation representations? To assess this, we examined the response among whisking-responsive neurons (**Figure S4A**). Whisking responses (mean whisking-onset aligned ΔF/F, Methods) among opsin-expressing neurons in both dual-stimulus induction mice (**Figure S4B**; from 0.08 ± 0.02 to 0.08 ± 0.03, P=1, Wilcoxon signed rank test comparing response before and after induction, N=7 mice) and photostimulation-only induction mice remained unchanged (from 0.05 ± 0.02 to 0.04 ± 0.02, P = 0.818, N=6 mice). Among opsin non-expressing neurons, whisking responses were also stable, both for dual-stimulus induction mice (from 0.09 ± 0.02 to 0.11 ± 0.04, P = 0.318) and photostimulation-only induction mice (from 0.08 ± 0.03 to 0.08 ± 0.03, P = 0.818). Thus, responsiveness declines are confined to touch and photostimulation responsive populations.

Did changes in opsin expression contribute to the observed changes in responsiveness? We used levels of mScarlet fluorescence (‘redness’; Methods) as a proxy for opsin expression (**Figure S5A**). We found that mean redness did not change following induction (**Figure S5B**), implying that opsin levels were stable. Moreover, pre-induction redness could reliably predict post-induction redness, implying that the opsin-expressing population was stable in membership (**Figure S5C, D**). If opsin levels remained unchanged yet photo-responsiveness declined, this may be due to a change in the degree to which opsin levels predicted response. We found that the correlation between mScarlet fluorescence, quantified as ‘redness’ (Methods) declined for both types of induction (**Figure S2D**).

We thus find that aggregate responsiveness among both touch and photostimulus responsive opsin-expressing neurons declines, consistent with dual-stimulus induction engaging homeostatic renormalization of experimentally elevated network activity (Gainey and Feldman, 2017). Moreover, the photostimulus responsive population exhibited greater membership stability than the touch responsive population.

### Touch-photostimulation training increases spontaneous activity correlations

Given that touch and dual-representation neurons exhibit higher within-group spontaneous correlations than purely photoresponsive neurons, we next asked if induction altered spontaneous activity correlations. Specifically, we were interested in whether microstimulation would enhance correlations within the photoresponsive population and between the photoresponsive and touch responsive populations, as this would be suggestive of enhanced connectivity within and between these populations (Cossell *et al*., 2015; Ko *et al*., 2011).

In individual mice, pairwise correlations during spontaneous activity among photostimulation responsive neurons increased for both opsin-expressing and opsin non-expressing neurons (**Figure 7A**). In opsin-expressing neurons, pairwise correlations among photostimulus-responsive neurons increased from 0.02 ± 0.01 to 0.06 ± 0.03 following dual-stimulus induction (P=0.016, Wilcoxon signed rank test comparing correlations pre- and post-induction, paired by animal; **Figure 7B**). Dual-stimulus induction also increased correlations among photostimulus-responsive opsin non-expressing neurons, from 0.03 ± 0.02 to 0.06 ± 0.02 (P=0.016). Photostimulation-only induction, however, did not produce a change in correlations among either opsin expressing (P=0.688) or opsin non-expressing (P=0.844) photostimulus-responsive neurons. Thus, dual-stimulus induction, but not photostimulation-only induction, increases correlations among photoresponsive neurons.

**Figure 7.**
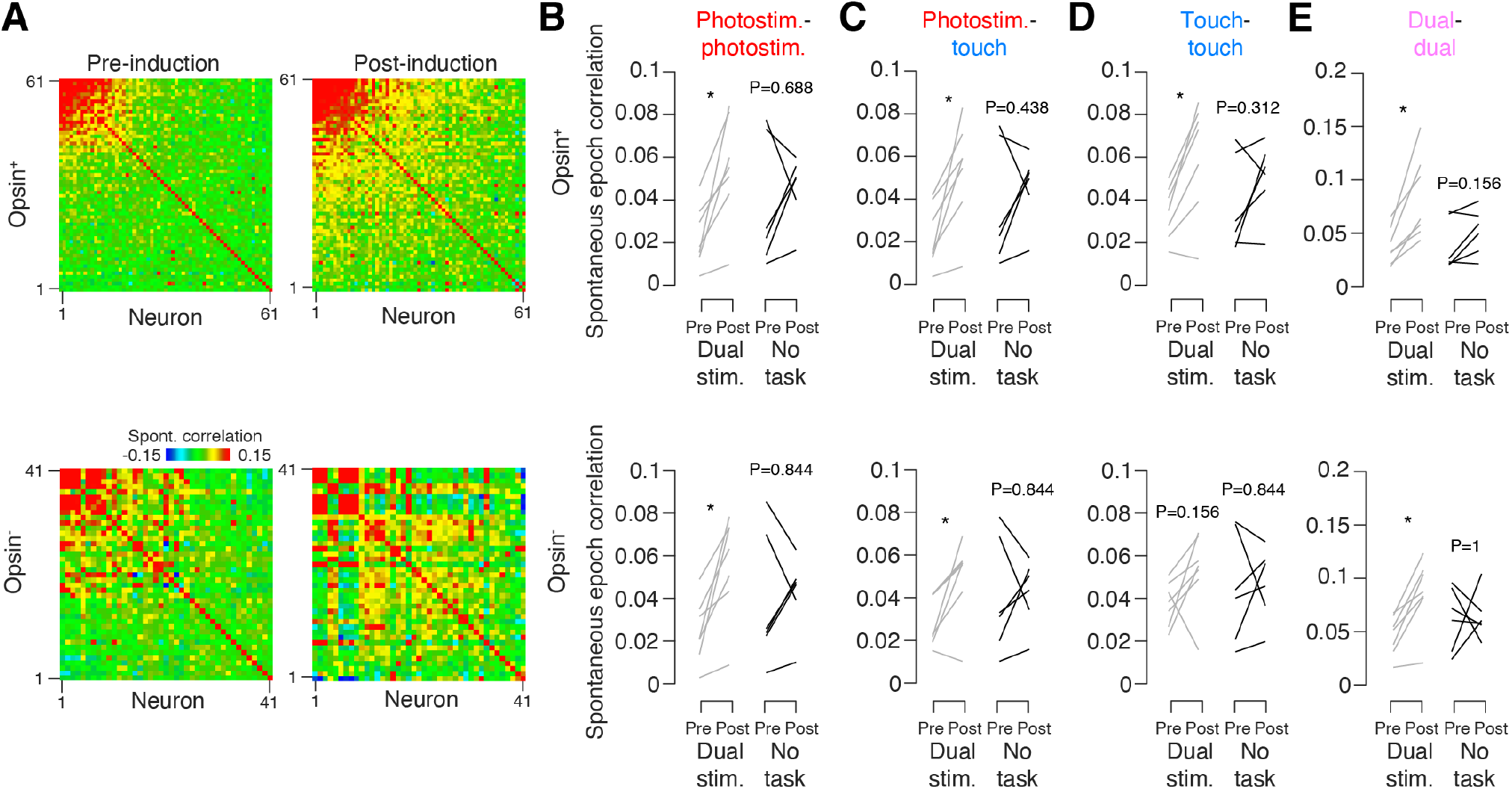
Spontaneous activity correlations before and after induction. (A) Spontaneous pairwise correlation matrices for the photostimulation-responsive population in an example mouse before (left) and after (right) dual-stimulus induction. Top, opsin-expressing population; bottom, opsin non-expressing. Neurons are sorted by their mean correlation to the other neurons within the population. (B) Mean pairwise correlations outside the stimulus epoch (‘spontaneous’ correlations; Methods) before and after induction among photoresponsive neurons. Grey, mice that were exposed to dual-stimulus induction (N=7); black, mice that were exposed to photostimulus-only induction (N=6). P-values provided for signed rank test comparing fraction before and after induction. ***, P < 0.001; **, P < 0.01 ; *, P < 0.05. Top, opsin-expressing neurons; bottom, opsin non-expressing neurons (C) As in (B), but for correlations between touch responsive and photoresponsive neurons. (D) As in (B), but for touch responsive neurons. (E) As in (B), but for dual-stimulus responsive neurons.

We next asked whether dual-stimulus induction changed the correlations between touch and photoresponsive populations. We found that for opsin-expressing neurons, dual-stimulus induction increased pairwise correlations between photoresponsive and touch responsive neurons from 0.02 ± 0.02 to 0.05 ± 0.02 (P=0.016; **Figure 7C**). Similarly, among opsin non-expressing neurons, dual-stimulus induction increased pairwise correlations between touch and photoresponsive neurons from 0.03 ± 0.01 to 0.05 ± 0.02 (P=0.016). Photostimulation-only induction did not alter photoresponsive-touch correlations for either opsin-expressing (P=0.438) or opsin non-expressing (P=0.844) neurons.

Dual-stimulus induction also increased correlations among opsin-expressing touch neurons, from 0.03 ± 0.01 to 0.06 ± 0.03 (P=0.031; **Figure 7D**), though touch-touch correlations remained unchanged among opsin non-expressing neurons (0.04 ± 0.01 to ± 0.02, P=0.156). Photostimulation-only induction did not alter touch-touch correlations among opsin-expressing (P=0.312) or opsin non-expressing neurons (P=0.844). Finally, dual-representation neurons showed the greatest increases in spontaneous correlations. Among opsin-expressing neurons, correlations increased from 0.04 ± 0.02 to 0.08 ± 0.04 (P=0.016; **Figure 7E**), and from 0.05 ± 0.02 to 0.09 ± 0.03 among opsin non-expressing neurons (P=0.016). Photostimulation-only induction again did not alter correlations (opsin-expressing, P=0.156; opsin non-expressing, P=1).

Thus, combined touch and photostimulation induction resulted in increased spontaneous activity correlations within photoresponsive, touch, and dual-representation populations, and between photoresponsive and touch responsive populations. Exposure to photostimulation-only induction did not alter spontaneous activity correlations in any population.

## DISCUSSION

We find that optogenetically stimulating a population of vS1 pyramidal neurons disproportionately engages existing whisker touch but not whisker movement neurons (**Figure 2**). Given that touch but not whisking vS1 neurons are recurrently coupled (Peron *et al*., 2020), our work suggests that cortical microstimulation engages recurrent pattern completion (Douglas and Martin, 2007). This is supported by our observation that spontaneous pairwise correlations among neurons that are both photoresponsive and touch responsive are comparable to correlations among touch neurons, whereas neurons that are purely photoresponsive show lower pairwise correlations (**Figure 3**). We further find that repeated co-activation of touch and photostimulus-responsive neurons increases representational overlap among these neurons (**Figure 4, 5**) and enhances spontaneous activity correlations within and between the touch and photostimulus responsive populations (**Figure 7**). Thus, we show that intrinsic network properties constrain externally introduced cortical activity and that this effect can be enhanced through repeated co-presentation of stimuli.

Whether activating a random subset of an area’s excitatory neurons should engage the same neurons as naturalistic stimuli is unclear. Cortical activity is typically confined to a small subspace of possible network activity patterns (Avitan and Stringer, 2022; Chung and Abbott, 2021; Tsodyks *et al*., 1999), potentially due to elevated recurrent connectivity (Koren and Deneve, 2017; Okun *et al*., 2015). Consistent with this, spontaneous activity resembles evoked activity in visual (Berkes et al., 2011; Kenet *et al*., 2003), auditory, and somatosensory cortices (Luczak *et al*., 2009). Moreover, local recurrent connections are highly specific with small subnetworks of highly and bidirectionally connected neurons embedded in a broader network with far lower connectivity (Perin *et al*., 2011; Song *et al*., 2005). These subnetworks likely exhibit similar sensory tuning, as neurons with similar tuning and higher pairwise correlations exhibit higher connectivity (Cossell *et al*., 2015; Harris and Mrsic-Flogel, 2013; Ko *et al*., 2011). This suggests that cortical recurrence should enable pattern completion (Douglas and Martin, 2007), a prediction borne out in experiments where direct activation of a small number of visual cortical neurons with particular tuning can drive activity in other neurons with similar tuning (Carrillo-Reid *et al*., 2019; Marshel *et al*., 2019). Our work suggests that even non-specific stimulation of cortex engages recurrence-mediated pattern completion which, in vS1, results in activation of existing touch but not whisking responsive populations. That opsin non-expressing neurons show greater overlap among photoresponsive and touch responsive neurons than opsin-expressing neurons bolsters this view, since opsin non-expressing neurons experience less direct drive from the photostimulus than opsin expressing neurons and should therefore be more constrained by intrinsic network properties.

In addition to excitatory recurrence, photostimulation of cortical neurons in vS1 L2/3 engages strong feedback inhibition within a few milliseconds that typically lasts up to ∼50 ms (Mateo et al., 2011). Moreover, though L2/3 pyramidal neurons with near-identical tuning amplify one another’s responses, the dominant interaction among neurons with even slightly different tuning is mutual suppression (Chettih and Harvey, 2019; Oldenburg et al., 2022). Such inhibition is likely to suppress responses, especially among downstream (i.e., opsin non-expressing) neurons. This suggests that our work provides a lower bound for overlap, and that actual engagement of existing representations could be enhanced either by mitigating inhibition or stimulating a far smaller number of neurons, reducing the recruitment of feedback inhibition.

Given the existence of a small subset of L2/3 vS1 neurons that are unusually touch responsive due to higher excitability (Crochet et al., 2011), it is possible that both natural and artificial excitation engages this subpopulation and that excitability, and not connectivity, account for the elevated overlap among photoresponsive and touch responsive populations. However, whisking neurons, which like touch neurons are part of the sparse and responsive minority of vS1 pyramidal neurons (Peron *et al*., 2015a), are not disproportionately engaged via optical microstimulation. Because touch, but not whisking, neurons exhibit recurrent amplification (Peron *et al*., 2020), the lack of elevated overlap with this population argues against excitability as the only explanation. Nevertheless, our experiments do not preclude this as a possible contributing mechanism, and it is likely that both excitability and connectivity play a role in explaining our results.

Repeated stimulation of small (Kim et al., 2016) and large (Carrillo-Reid et al., 2016; Zhang et al., 2020) cortical populations increases responsiveness among the stimulated neurons due to elevated connectivity, increased excitability (Alejandre-García et al., 2020), or both. Our observation of declining overall responsiveness (**Figure 6**) suggests that our paradigm does not enhance excitability, implying enhanced connectivity as the dominant mechanism. This is consistent with the plasticity rules governing excitatory-excitatory synapses within L2/3 of vS1 (Banerjee *et al*., 2014), as well as simulations showing that repeated co-activation of neurons drives enhanced connectivity (Clopath *et al*., 2010; Litwin-Kumar and Doiron, 2014). Nevertheless, further experiments are needed to determine the precise mechanism driving increased overlap.

Sensory cortical feedback is important for the design of effective neural prosthetics (Pandarinath and Bensmaia, 2021; Tyler, 2015). Such feedback can reproduce subtle features of somatosensory percepts, such as vibration frequency (Romo *et al*., 1998), and recapitulate tactile sensations that improve the control of prosthetic limbs (Flesher et al., 2021). The elevated overlap between photoresponsive and touch responsive populations suggests that even non-specific cortical feedback may evoke naturalistic touch-like activity patterns and thus, touch sensations. This may explain the effectiveness with which stimulation of somatosensory cortex can perceptually bias animals trained to use natural somatosensory input (O’Connor et al., 2013; Romo *et al*., 1998), and the longstanding observation that cortical stimulation can evoke both naturalistic percepts (Penfield and Boldery, 1937) and movements (Graziano *et al*., 2002). Indeed, our work suggests that direct cortical stimulation may be able to evoke naturally occurring patterns of activity without sophisticated cellular-resolution stimulation approaches. Moreover, our induction experiment suggests that appropriate induction paradigms should enhance the ability of microstimulation to engage naturalistic activity patterns.

We exposed mice either to a rewarded task with paired photostimulation and touch, or reward-free photostimulation, with the paired task producing enhanced overlap between touch and photoresponsive populations. It thus remains unclear whether enhanced overlap depends on co-stimulation of the touch and photoresponsive neurons, on the animal being engaged due to the presence of reward, or both. Cortical plasticity is often dependent on task engagement: in auditory cortex, enhanced sensitivity to a repeatedly presented tone depends on reward and hence engagement (Kuchibhotla et al., 2017; Weinberger, 2004). In a separate study using a similar paradigm to the one presented here in which mice were trained to discriminate the number of optogenetic pulses delivered to vS1 for a water reward (Pancholi et al., 2022), overall responsiveness declined following training, analogous to our dual-stimulus induction mice but unlike the photostimulus-only induction mice. If photostimulation in a rewarded context is sufficient to drive enhanced overlap, it would suggest an even easier strategy for generating elevated overlap, an approach that could benefit the design of somatosensory feedback prostheses.

In sum, we show that optical microstimulation engages natural sensory representations, but only those whose constituent neurons are known to be recurrently coupled. Moreover, repeated co-presentation of photostimulation and natural sensory input increases the degree to which photostimulation engages the natural sensory representation. Our work thus shows that inherent network structure constrains cortical activity, and that even untargeted direct cortical stimulation can engage existing cortical subnetworks.

## ACKNOWLEDGMENTS

We thank Robert Froemke, Anthony Movshon, Michael Long, and members of the Peron lab for discussion. This work was supported by the Whitehall Foundation and the National Institutes of Health (R01NS117536; F31NS120483; T32GM007308).

## AUTHOR CONTRIBUTIONS

R.P. and S.P. designed the study. R.P. and A.S-Y. carried out all the experiments. R.P. and S.P. performed data analysis and wrote the paper.

## FIGURE LEGENDS

**Figure S1.**
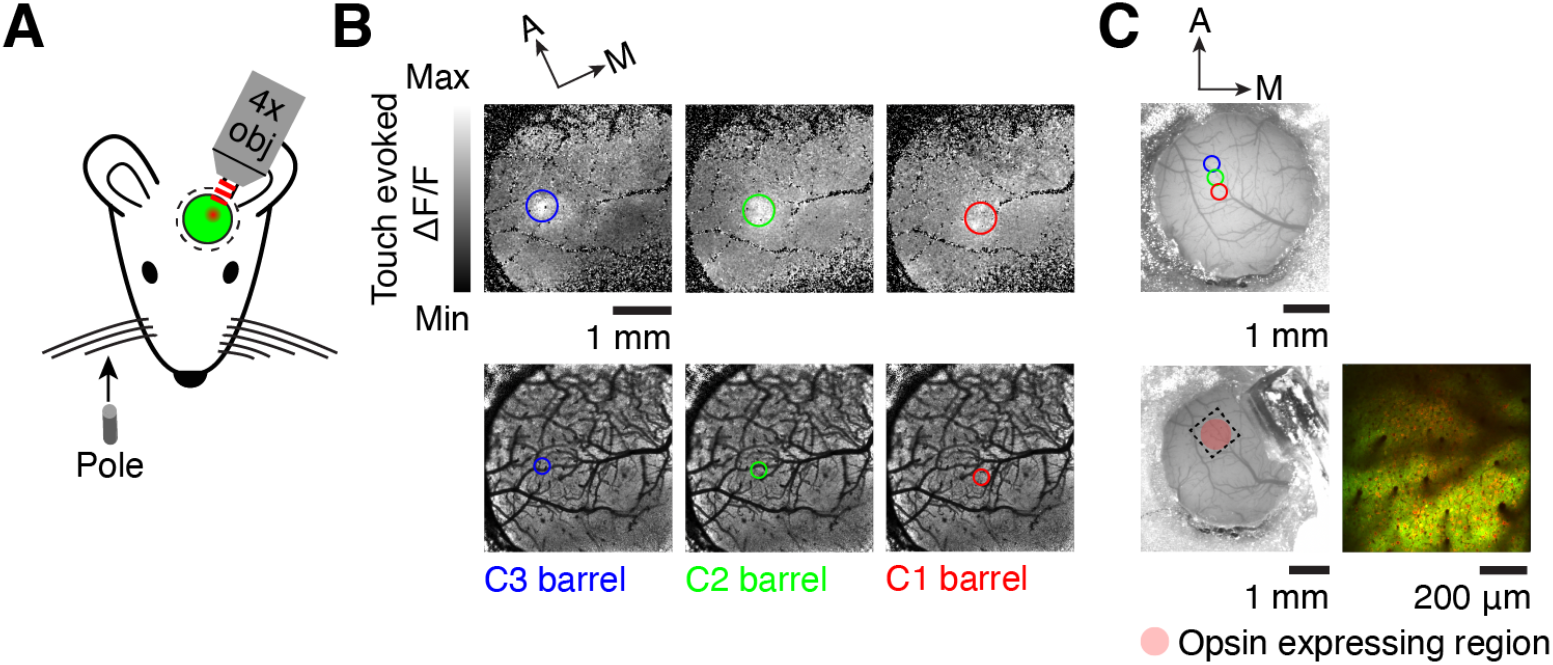
Targeting single barrel for superficial opsin expression. (A) Barrel mapping was conducted by moving a pole toward the C1, C2, or C3 whiskers during widefield (4x) two-photon calcium imaging. (B) Touch-evoked ΔF/F following whisker stimulation. ΔF/F after touch by indicated whisker (top) and barrel column locations on corresponding vasculature (bottom). (C) Cranial window before (top) and after (bottom) LED placement. Higher magnification two-photon image of opsin-expressing area (right). Colored circles indicate the identified barrel locations from (B), and the dotted circle indicates the site of opsin expression.

**Figure S2.**
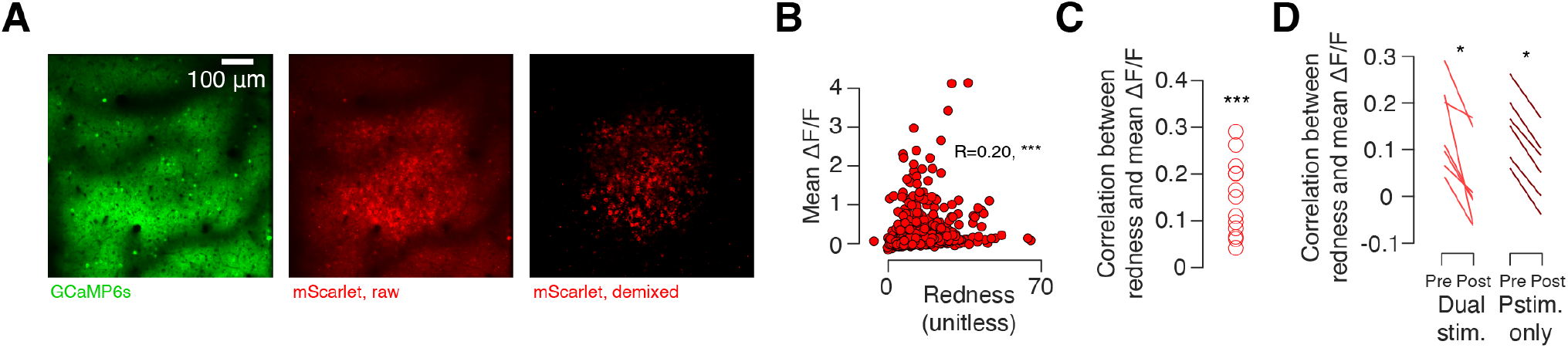
Relationship between opsin expression and response to photostimulation. (A) Opsin expression was assessed using the linearly demixed red PMT channel image (Methods). Left, raw GCaMP6s image from the green PMT channel. Middle, raw mScarlet image from the red PMT channel. Right, linearly demixed mScarlet image – ‘redness’. (B) Relationship between redness of individual opsin-expressing neurons and the mean evoked ΔF/F following photostimulation. P-value indicated for Pearson correlation between redness and evoked ΔF/F. ***, P < 0.001; **, P < 0.01 ; *, P < 0.05. (C) Pearson correlation between redness and evoked ΔF/F for all animals (N=13). P-value indicated for Wilcoxon signed rank test comparing the measured median to 0. (D) Pre- and post-induction Pearson correlation between redness and evoked ΔF/F. Left, dual-stimulus induction. Right, photostimulation-only induction. P-value indicated for Wilcoxon signed rank test comparing pre vs. post-induction correlation, paired by animal.

**Figure S3.**
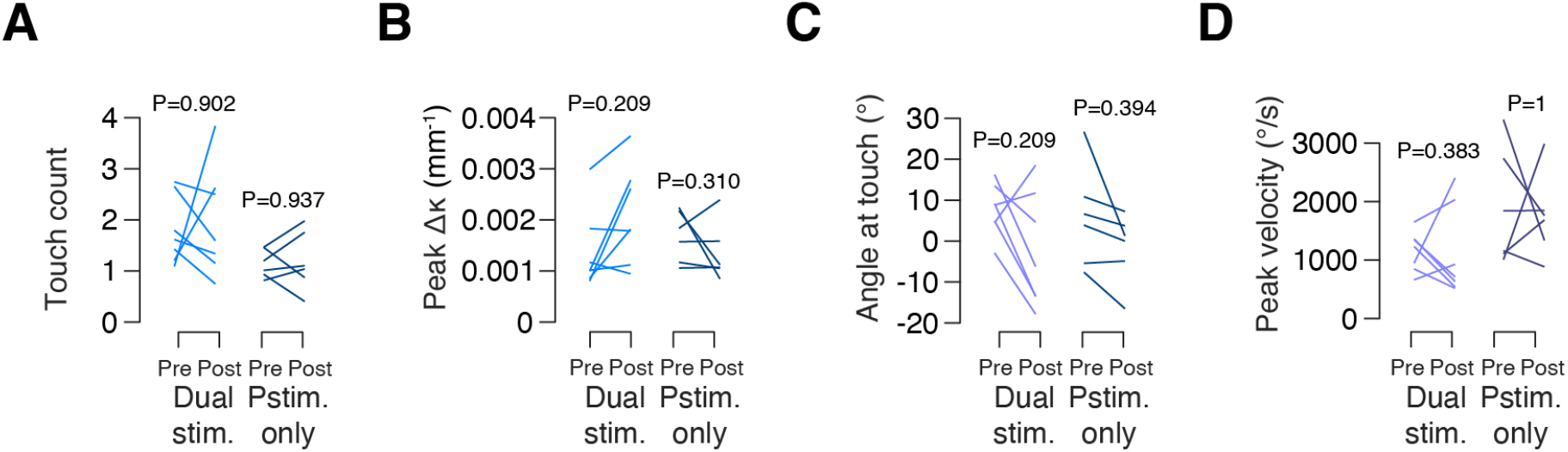
Vibrissal kinematics pre- and post-induction. (A) Number of touches before and after induction, averaged across all trials in a session. Left, dual-stimulus induction. Right, photostimulation-only induction. P-value indicated for Wilcoxon signed rank test comparing pre vs. post-induction value, paired by animal. (B) As in A, but for the peak curvature change (Δκ) experienced during a trial, averaged across trials. (C) As in A, but showing the mean angle of the whisker at touch for a trial, averaged across trials. (D) As in A, but showing the peak angular velocity during the trial, averaged across trials.

**Figure S4.**
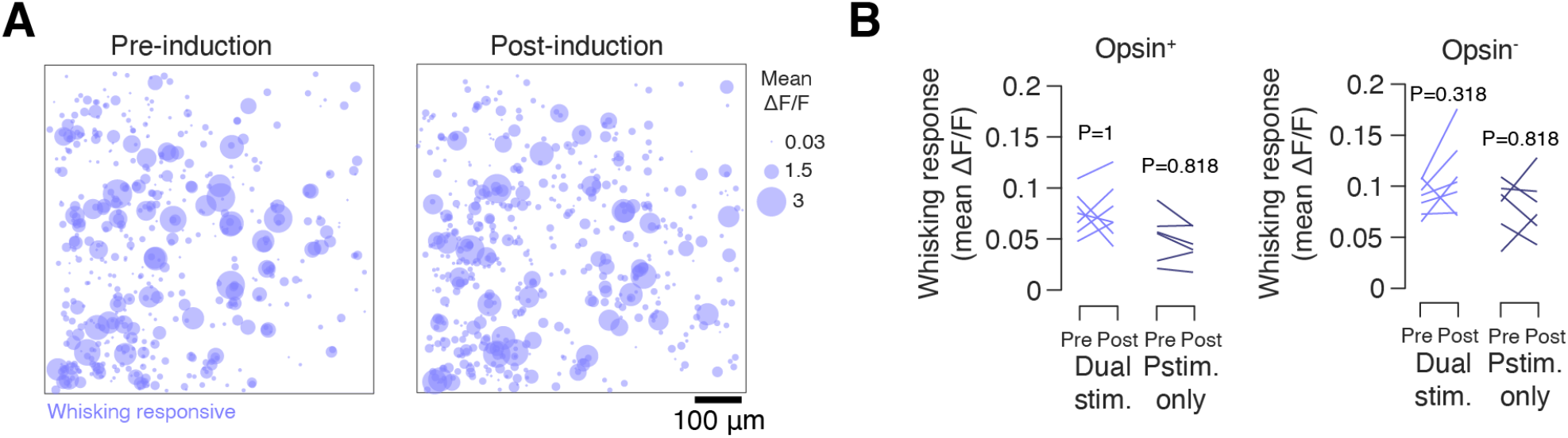
Change in whisking populations following induction. (A) Whisking population in an example mouse before (left) and after (right) dual-stimulus induction. Neurons are collapsed across six planes spaced 20 μm apart. Colored circles show the mean whisking-evoked ΔF/F. (B) Change in mean whisking-evoked ΔF/F among the top 10% of most whisking-responsive neurons following induction. Left, opsin-expressing; right, opsin non-expressing. In each case, we examined mice trained on dual-stimulus induction (N=7; lighter color), and mice that were repeatedly photostimulated with no task (N=6; darker color). P-values provided for signed rank test comparing response before and after induction. ***, P < 0.001; **, P < 0.01 ; *, P < 0.05.

**Figure S5.**
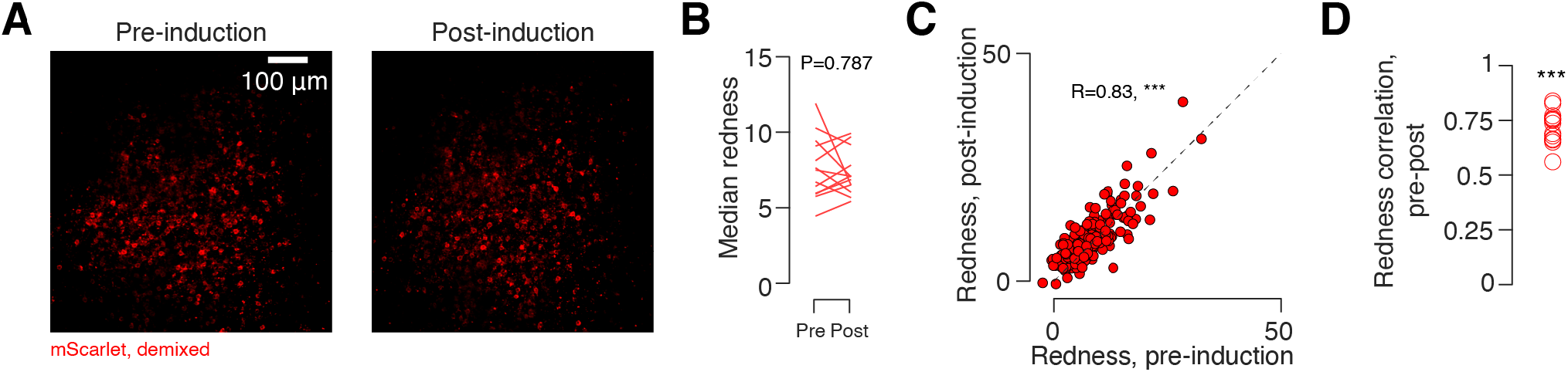
Stability of opsin expression. (A) Linearly demixed mScarlet image on the first and final days of induction in an example mouse. (B) Median redness before and after induction. P-value for Wilcoxon signed rank test comparing pre- vs. post-induction redness, paired by animal (N=13). ***, P < 0.001; **, P < 0.01 ; *, P < 0.05. (C) Pearson correlation between redness pre- vs. post-induction in an example animal. P-value given for Pearson correlation. (D) Pearson correlation between pre- and post-induction redness for all animals. P-value indicated for Wilcoxon signed rank test comparing pre vs. post-induction correlation to 0, paired by animal.

## METHODS

### Mouse lines

We used adult male (n = 9) and female (n = 4) Ai162 (JAX 031562) X Slc17a7-Cre (JAX X 023527) (Daigle *et al*., 2018) mice throughout this study (**Table S1**). These mice express GCaMP6s exclusively in excitatory neurons throughout cortex in a tetracycline transactivator-dependent manner. To suppress expression during development, breeders were fed doxycycline chow (625 mg/kg doxycycline; Teklad), and all pups therefore received doxycycline until weaning. All animal procedures complied with protocols approved by New York University’s University Animal Welfare Committee.

### Surgical preparation

Mice (6-9 weeks old) were anesthetized with isoflurane during viral injections, window implantation, LED placement, and whisker trimming (3% induction, 1.4% maintenance). During surgery, a titanium headbar was affixed to the skull with cyanoacrylate (Vetbond). A circular craniotomy (3.5 mm diameter) was then drilled over left vS1 (center: 3.3 mm lateral, 1.7 mm posterior from bregma) using a dental drill (Midwest Tradition, FG 1/4 drill bit).

Following removal of the bone flap, a viral vector encoding the soma-localized opsin ChRmine and the red fluorophore mScarlet (AAV-8-CaMKIIa-ChRmine-mScarlet-Kv2.1-WPRE, 2.48×10^13^ vg/mL, diluted either 1:200 or 1:500 in 1X PBS; generously provided by Karl Deisseroth) was injected into vS1. A glass capillary (Wiretrol II, Drummond) was pulled to have a tip diameter of 25 μm using a micropipette puller (P-97, Sutter Instrument) and then beveled to a 30° tip. The pipette was back-filled with mineral oil (M5904, Sigma-Aldrich) and 2 μL of viral solution drawn into the tip using a plunger. One, 20 nL injection or three, 100 nL injections were made 250 μm below the dura (**Table S1**). When performing multiple injections, injections were spaced 400 μm apart in a triangle centered on the craniotomy. For each injection: (1) the pipette was lowered into the brain at a rate of 300 μm/min, (2) there was a 1-minute pause, (3) virus was injected at a rate of 20 nL/min using a hydraulic micromanipulator (Narishige MO-10), (4) there was a 2 minute pause, and (5) the pipette was withdrawn at a rate of 300 μm/min with (6) an additional 1 minute pause at a depth of 125 μm below the dura. Following injections, the dura was removed using a pair of fine forceps (Fine Science Tools). A double-layer cranial window (4.5 mm external diameter, 3.5 mm inner diameter; #1.5 coverslip; adhered with Norland 61 UV glue) was then placed over the craniotomy and the skull covered with dental acrylic (Orthojet, Lang Dental) to affix the cranial window and headbar. Mice were post-operatively injected with 1 mg/kg of buprenorphine SR and 5 mg/kg of ketoprofen.

Following surgical recovery, mice were trimmed to whiskers C1-3 and placed on water restriction. To confirm that the area of opsin-expression fell within vS1, the location of the C1-3 barrel columns was identified (**Figures S1A**-**C**). Mice were first head fixed and neural activity throughout the cranial window observed at coarse resolution (4X objective; 3 × 3 mm field of view). A pole was brought into the whisking plane and moved in a posterior direction (1-2 mm) so as to touch each whisker individually. The touch-evoked ΔF/F was used to localize the whisker barrel column.

Following barrel identification, the photostimulation LED was placed. LEDs (590 nm, LXZ1-PL01, Lumileds) were fabricated by soldering 3 cm of polyurethane enameled copper wire (34 AWG) to each pad of the LED. A 2-pin, flat flex cable connector (Digikey) was soldered to the free ends of the copper wire and secured with epoxy (Devcon). The LED was waterproofed using a thin layer of clear nail polish (Sally Hansen). LEDs were affixed to the animals’ cranial windows under anesthesia. The anterior medial edge of the craniotomy was exposed by drilling away a 60° arc of dental cement and covering the drilled area with cyanoacrylate. Electrical tape was placed over the headbar to prevent current from passing between the LED wires and the headbar. The LED was placed over the drilled area 0.5-1 mm away from the site of opsin expression and at a 30° angle relative to the plane of the window (**Figure 1A**). The LED and copper wires were secured to the dental cement using cyanoacrylate, and the LED connector was secured to the posterior edge of the headbar in a similar manner. Waterproofing was confirmed by placing water on the cranial window and ensuring that no current passed between the water and the LED.

### Photostimulation system

Optogenetic stimulus delivery was controlled by a LabJack T7 driven by a Raspberry Pi. Stimulus voltage waveforms were generated using custom MATLAB software on a separate computer and sent to the Raspberry Pi. The Raspberry Pi loaded waveforms onto the LabJack and waited for a stimulus trigger. Upon receipt of the trigger, the waveforms were sent to the stimulation and masking flash LED drivers (T-cube, ThorLabs) and the PMT shutters. The signal to the stimulation LED was terminated using a 4-pin, flat flex cable connector (Digikey) that could be mated to the LED connector when the animal was head-fixed. The masking flash consisted of 3 LEDs (595 nm, XPEBAM-L1-0000-00A01, Cree LED) that were spectrally matched to the stimulation LED and placed around the animal’s face to illuminate the eyes.

### Two-photon microscopy

Calcium imaging was performed using a custom two-photon microscope (http://openwiki.janelia.org/wiki/display/shareddesigns/MIMMS). The microscope consisted of a 940 nm laser (Chameleon Ultra 2, Coherent), a Pockels cell (350-80-02, Conoptics), two galvanometer scanners (6SD11268, Cambridge Technology), a resonant scanner (6SC08KA040-02Y, Cambridge Technology), a 16x objective (N16XLWD-PF, Nikon), an emission filter for green fluorescence (FF01-510/84-30, Semrock), an emission filter for red fluorescence (FF01-650/60, Semrock), and two GaAsP PMTs (H10770PB-40, Hamamatsu) and two PMT shutters (VS14S1T1, Vincent Associates).

Imaging data was acquired using Scanimage (Vidrio Technologies). Three 800-by-800 μm imaging planes axially spaced 20 μm apart were acquired at a rate of ∼7 Hz (one subvolume). Two continuous subvolumes were acquired for each animal and alternated every ∼100-150 trials. The objective was moved axially with a piezo (P-725KHDS, Physik Instrumente). Power was depth-adjusted by the acquisition software with an exponential length constant of ∼250 μm. Imaging data were processed on the NYU High Performance Computing cluster using a semi-automated software pipeline (Peron *et al*., 2015b). The pipeline included image registration, segmentation, neuropil subtraction, ΔF/F computation, and calcium event detection. Imaging planes for each day were aligned using motion-corrected mean images generated from the first imaging session (Huber et al., 2012).

Opsin expression was measured using mScarlet fluorescence collected in the red PMT channel. For each pixel, we calculated a ‘redness score’, which consisted of the red channel pixel value following linear de-mixing to remove cross-talk introduced by GCaMP fluorescence. For each neuron, an overall redness score was found by computing the mean de-mixed redness across its pixels. For each mouse, we manually selected a value above which neurons were considered opsin-expressing. Neurons with a redness below a second, lower threshold were considered opsin non-expressing. Neurons with an intermediate redness were considered ambiguous and were excluded from population-specific analyses.

### Identification of touch and whisking neurons

An imaging session was used to assess touch and whisking sensitivity prior to the first induction session. During this session, a 0.5 mm diameter pole was brought into the whisking plane for 1 s (**Figure 1B**). To encourage whisking, mice were rewarded with water following a 0.5 s delay period. Water was delivered from a single lickport if mice licked within 2 s of the end of the delay period (‘response’ epoch). Mice were given a fixed amount of time (1-2 s) to collect water upon responding. Trials lasted 10-12 s, with variability resulting from when the response occurred during the response epoch. High-speed whisker videography was used to track whiskers throughout the imaging session and classify neurons as either touch- or whisking-responsive (see below).

### Induction protocols

Animals were presented with one of two induction protocols: either photostimulus-only or dual photostimulus and touch (**Table S1**). For mice exposed to the photostimulation-only induction protocol, mice were presented with 9 pulses (5 ms duration, 20 Hz frequency) from the photostimulation LED on 50% of trials. During the photostimulation epoch, the PMT shutters were closed and the masking flash LED was illuminated (9 pulses, 15 ms duration, 20 Hz frequency, pulses centered on the photostimulation LED pulses). On the remaining 50% of trials, no stimulus was presented but the masking flash and PMT shutters were still activated. This allowed us to assess whether neurons were visually responsive or responsive to the auditory cue produced by the shutter. Trials proceeded with an interstimulus interval of 10 s. Mice were presented with 453.1 ± 33.2 trials per session for 10 sessions.

For mice exposed to dual photostimulus and touch induction, four trial types were used. On 50% of trials, no stimulus was presented but the masking flash and PMT shutters were activated. This allowed us to assess whether neurons were visually responsive or responsive to the auditory cue produced by the shutter. On 16.67% of trials, only a photostimulus was presented; on another 16.67% of trials, only a pole stimulus was presented; on the remaining 16.67% of trials, both stimuli were presented (**Figure 4A**). Mice were presented with 387.1 ± 54.8 trials per session for 10 sessions. In the dual-stimulus induction mice, to encourage whisking, mice were given a water reward for licking the right of two lickports on trials when a stimulus was present, and the left of two lickports when stimuli were absent. Stimulus presentation and response were separated by a 0.5 s delay.

### Whisker videography and vibrissal representation analysis

Mice were trimmed to the C1-2 or C2-3 whiskers (**Table S1**) prior to the pre-induction touch and whisking sensitivity assessment and regularly trimmed over subsequent days. Whisker video was acquired from a CMOS camera (Ace-Python 500, Basler) running at 400 Hz and 640 × 352 pixels with a telecentric lens (TitanTL, Edmund Optics). A pulsed 940 nm LED (SL162, Advanced Illumination) was used to illuminate the camera’s field-of-view (typical exposure and illumination duration: 200 μs). Custom MATLAB software was used to control video acquisition. 7 s of each trial were recorded, including 1s prior to pole movement, the period when the pole was in reach, and several seconds after the pole was retracted.

Whisker video was processed on NYU’s High Performance Computing (HPC) cluster. Candidate whiskers were first detected using the Janelia Whisker Tracker (Clack et al., 2012). Whisker identity was then refined and assessed across a single session using custom MATLAB software (Peron *et al*., 2020; Peron *et al*., 2015b). Following whisker assignment, whisker curvature (κ) and angle (θ) were calculated at specific locations along the whisker’s length. Whisking setpoint, amplitude, and velocity were computed by decomposing the whisker angle (θ) using the Hilbert transform (Kleinfeld and Deschenes, 2011). Whisker bout onset was defined as the point where the whisking amplitude reached at least 10°. Change in curvature, Δκ, was calculated relative to a baseline, angle-dependent curvature value obtained during periods when the pole was out of reach. Following automatic touch detection, touch assignment was manually curated using a custom MATLAB user interface (Peron *et al*., 2015b). As per convention, protractions were assigned negative Δκ values.

Neurons were assigned to the touch or whisking representations based on the evoked ΔF/F response following touch or whisking bout onset. First, a baseline ΔF/F was computed for each neuron by taking the mean ΔF/F over the 5.5 s prior to each touch or whisking bout onset. Then, a response ΔF/F was computed by taking the mean ΔF/F over the 2 s following touch or whisking bout onset. The evoked ΔF/F was the difference between the response and control ΔF/F. The mean evoked ΔF/F was then computed across trials for each neuron. Neurons were considered to be ‘whisking’ neurons if their mean evoked ΔF/F fell into the top 10% of neurons following whisking bout onset. Similarly, neurons were considered ‘touch’ neurons if their mean evoked ΔF/F fell into the top 10% of neurons following touch onset. This cut-off is in line with estimates of the percentage of touch (12%) and whisking (12%) L2/3 excitatory neurons in the whisker barrel column (Peron et al. 2015).

### Photoresponsive representation analysis

Neurons were classified as responsive or non-responsive by computing the photostimulation-evoked ΔF/F. A baseline ΔF/F was computed for each neuron by taking the mean of the ∼5.5 s (39 frames) preceding shutter closure on each trial. The post-stimulation ΔF/F was calculated as the mean ΔF/F of the two frames immediately following shutter reopening. For each trial, the photostimulation-evoked ΔF/F was found by taking the difference between the post-stimulation ΔF/F and the baseline ΔF/F. Neurons were considered photoresponsive in any given session if they fell within the top 10% of neurons based on the mean photostimulation-evoked ΔF/F computed across photostimulation trials.

### Overlap analysis

We defined each stimulus representation as the top 10% of neurons responsive to that stimulus (see vibrissal representation analysis and photoresponsive representation analysis above). Neurons were selected from the same population, either opsin-expressing or opsin non-expressing. For example, consider animal an015145 (**Supplementary Table 1**). This animal had 724 opsin-expressing neurons and 3304 opsin non-expressing neurons. Therefore, the touch representation was defined as the 72 opsin-expressing and 330 opsin non-expressing neurons with the greatest mean touch-evoked ΔF/F. This process was repeated for the whisking and photoresponsive representations using the mean whisking-evoked ΔF/F and photostimulus-evoked ΔF/F, respectively. Because the size of each representation was fixed at 10% of the recorded population size, the overlap between two representations expected by chance was fixed at 1%. For animal an015145, this was 7.24 opsin-expressing and 33.04 opsin non-expressing neurons. The normalized overlap was then computed by dividing the actual number of neurons shared between two representations by the number expected by chance. A two-tailed Wilcoxon signed rank test was used to assess if this value was significantly different from one.

### Spontaneous activity correlation analysis

To compute pairwise correlations between neurons during non-stimulus epochs, we restricted our analysis to the touch-only sessions. We sampled from the time period 5 s after the pole stimulus was removed and computed correlations between ΔF/F vectors for all pairs of simultaneously imaged neurons.

## Statistical analysis

For comparisons between paired samples, we used the Wilcoxon signed rank test. For unpaired samples, we used the Wilcoxon rank sum test. For correlation tests, Pearson’s correlation was used to identify a linear correlation coefficient (R) and test for significance.

## SUPPLEMENTARY MATERIALS

**Supplementary Table 1.** List of animals. All mice were transgenic Ai162 X Slc17a7-Cre (Daigle *et al*., 2018), expressing GCaMP6s exclusively in excitatory neurons. All mice except an014261 were injected with 3, 100 nL injections using a 1:500 dilution; an014261 received 1, 20 nL injection with a 1:200 dilution.

| Animal ID | Sex | Age at training start (d) | Opsin<sup>+</sup> cell count | Opsin<sup>-</sup> cell count | Induction protocol |
| --- | --- | --- | --- | --- | --- |
| an014261 | M | 97 | 1094 | 3461 | Dual-stimulus |
| an014347 | F | 78 | 615 | 2950 | Dual-stimulus |
| an014350 | F | 75 | 589 | 3529 | Dual-stimulus |
| an015145 | M | 103 | 724 | 3304 | Dual-stimulus |
| an014317 | M | 114 | 733 | 2358 | Dual-stimulus |
| an016655 | M | 130 | 666 | 3652 | Dual-stimulus |
| an017772 | M | 83 | 828 | 2257 | Dual-stimulus |
| an014348 | F | 80 | 319 | 3002 | Photostimulation-only |
| an015742 | M | 82 | 391 | 2137 | Photostimulation-only |
| an015741 | M | 94 | 770 | 3165 | Photostimulation-only |
| an016663 | M | 94 | 576 | 2363 | Photostimulation-only |
| an015143 | M | 120 | 611 | 3145 | Photostimulation-only |
| an014349 | F | 114 | 1012 | 2315 | Photostimulation-only |

